# Systematic identification of cargo-carrying genetic elements reveals new dimensions of eukaryotic diversity

**DOI:** 10.1101/2023.10.24.563810

**Authors:** Emile Gluck-Thaler, Aaron A. Vogan

## Abstract

Cargo-carrying mobile elements (CCEs) are genetic entities that transpose diverse protein coding sequences. Although common in bacteria, we know little about the biology of eukaryotic CCEs because no appropriate tools exist for their annotation. For example, *Starships* are fungal CCEs whose functions are largely unknown because they require time-intensive manual curation. To address this knowledge gap, we developed starfish, a computational workflow for high-throughput eukaryotic CCE annotation. We applied starfish to 2, 899 genomes of 1, 649 fungal species and found that starfish recovers known *Starships* with >95% precision and accuracy while expanding the number of annotated elements ten-fold. Extant *Starship* diversity is partitioned into 11 families that differ in their enrichment patterns across fungal classes. *Starship* cargo changes rapidly such that elements from the same family differ substantially in their functional repertoires, which are predicted to contribute to diverse biological processes such as metabolism. Many elements have convergently evolved to insert into 5S rDNA and AT-rich sequence while others integrate into random locations, revealing both specialist and generalist strategies for persistence. Our work establishes a framework for advancing mobile element biology and provides the means to investigate an emerging dimension of eukaryotic genetic diversity, that of genomes within genomes.

## Introduction

Mobile elements are genetic entities found within organismal and organellar genomes that have evolved mechanisms to replicate themselves. Once dismissed as little more than genomic parasites, mobile elements are increasingly recognized to interact with each other and their host genomes in complex ways that lead to surprising outcomes (1–6). Although many mobile elements are expected to be evolving neutrally through genetic drift or to be under strong purifying selection (7–9), even short bursts of their activity can have drastic effects on their hosts’ evolutionary trajectories. For example, mobile elements have variously been associated with both local and global changes in host gene expression, with the spread of epigenetic marks, with the generation of loss-of-function mutants, and with genomic rearrangements facilitated through non-homologous ectopic recombination (10–14). Mobile element replication, which maximizes mobile element fitness, may result in intragenomic conflict if it simultaneously decreases host fitness, or intragenomic cooperation if instead it augments host fitness (15, 16). Similar to macroscopic ecosystems where species interactions beget emergent ecological and evolutionary outcomes, genomes resemble ecological arenas where interactions between different agents shape the trajectory of organismal ecology and evolution (17–19). Characterizing the community of genetic entities within an organism and their myriad interactions is therefore of central importance for deciphering the genetic bases of adaptation (20).

Mobile elements exist in a variety of forms and encode diverse strategies for their survival (21, 22). While many smaller elements like transposons consist solely of loci required for their own excision, integration and replication, others have evolved the ability to carry long stretches of DNA embedded within their boundaries. These cargo carrying elements (CCEs) are common features of prokaryotic genomes and often contain loci contributing to adaptive phenotypes (23–25). For example, integrative and conjugative elements (ICEs) and plasmids are key players in the evolution and dispersal of many important traits such as antibiotic resistance, heavy metal tolerance, and specialized metabolic pathways mediating both pathogenic and mutualistic species interactions (26–29). Through reshuffling genes amongst each other and their genomic hosts and through driving gene gain and loss, bacterial CCEs are important contributors to the generation of selectable genetic variation and the emergence of novel genotypes (22, 30).

Once thought to possess only simpler and smaller elements, eukaryotic genomes are proving to be rich and novel sources of large CCEs. The past several years have seen first reports of CCEs in diverse eukaryotic taxa ranging from fish to nematodes and algae (31–33) and the expectation is that the repertoire of eukaryotic CCEs will only continue to grow with the advent of population-level long-read sequencing (34). We recently described the *Starships*, a new group of CCEs mobilized by tyrosine recombinases (YRs) that are endemic in the largest kingdom of microbial eukaryotes, the fungi (35). *Starships* not only carry genes for their own mobilization but also carry diverse sets of accessory cargo with notable impacts on host phenotypes. The *Starship Enterprise* carries a meiotic gene drive that fuels the killing of spores that lack it (36); *Starship HEPHAESTUS* confers the ability to withstand heavy metal toxicity through carrying multiple genetic modules (37, 38); *Starship Horizon* carries the ToxA gene, enabling the plant pathogen *Pyrenophora tritici-repentis* to infect wheat plants (35, 39). *Starships* likely share certain properties with integrative bacterial CCEs like ICEs, such as the ability to transfer genes between individuals, to recombine and reshuffle genes amongst themselves, and are expected to be present in low copy numbers (22, 23, 35, 38, 40). However, much about *Starship* biology remains unknown, including their strategies for persistence and their impact on the evolution of key host phenotypes. Studying eukaryotic CCEs, like *Starships,* thus stands to not only reveal new rules governing CCE evolution but to shed light more broadly on the mechanisms of eukaryotic adaptation.

Assessing the impact of eukaryotic CCEs remains challenging due to the simple fact that it is difficult to determine if they are present within a genome. Many mobile element annotation tools are premised on the assumption that eukaryotic elements exist in multi-copy states, and therefore focus on detecting repetitive sequences instead of low copy number elements (41, 42). Other tools developed specifically for bacterial genomes predict CCEs based on the presence of known “core” genes that are commonly present among different elements (23). While such tools are useful for annotating CCEs with well-defined integration, excision, conjugation and regulatory modules, the sequence content of eukaryotic CCEs is largely unknown, precluding content-based predictions. As a result, reliable CCE annotation in eukaryotes is currently only possible through time-intensive manual curation, which imposes a significant bottleneck for high-throughput comparative evolutionary and functional studies.

To overcome this methodological gap, we developed starfish, the STARship FInder SHell, a computational workflow for CCE annotation that operates independently of element copy number and sequence content. As a proof of concept, we used starfish to annotate *Starship* elements in 2, 899 genomes from 1, 649 species sampled from all major families across the fungal tree of life. We used the resulting output to develop a phylogenetic framework for classifying *Starship* diversity and applied it to gain insight in the mode and tempo of eukaryotic CCE evolution.

## Materials and Methods

Starfish is a computational workflow for *de novo* giant mobile element annotation. It is organized into three modules that identify the coordinates of genes of interest (Gene Finder), the coordinates of elements associated with those genes (Element Finder), and the coordinates of larger genomic regions containing those elements (Region Finder) (Figure 1). The minimum inputs required to run starfish are two genome assemblies from different individuals. It is possible but not recommended to use individuals from different species due to challenges associated with detecting homologous sequences across distantly related species through nucleotide alignments. Although not necessary, the quality of starfish output is enhanced by providing predicted gene annotations, repeat annotations, gene orthogroup assignments, and as many genome assemblies from the same species as possible. Although we built starfish to help find *Starship* elements in fungal genomes, starfish can be used to find any integrative mobile element that shares the same basic architecture as a *Starship*: a “captain” gene with zero or more “cargo” genes downstream of its 3’ end. Auxiliary commands for visualizing output are also distributed as part of the workflow, enabling the generation of graphics for publication and visual quality control (43, 44). Also included are several commands that facilitate comparative analyses of interest, such as alignment and short read mapping searches for known elements. A step-by-step walkthrough as well as a manual describing the options and requirements for specific commands is available at the starfish wiki: https://github.com/egluckthaler/starfish/wiki/. All analyses presented here were conducted with starfish v1.0.0 (DOI: <u>10.5281/zenodo.8345762</u>)

**Figure 1:**
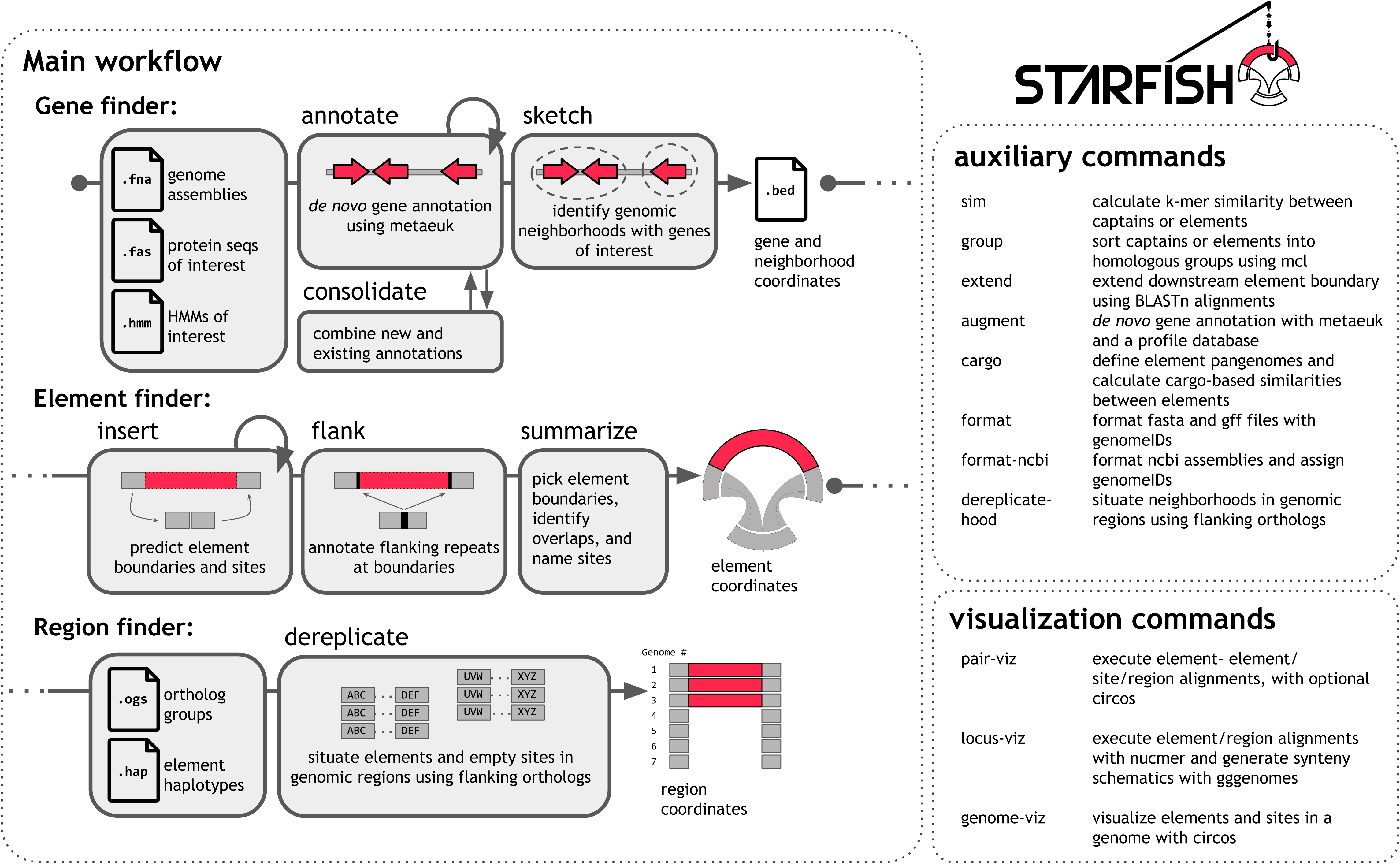
The starfish workflow and auxiliary commands. Starfish is organized into three modules: the Gene Finder Module uses metaeuk to de novo annotate genes of interest, typically *Starship* tyrosine recombinases. The Element Finder Module annotates *Starship* element coordinates based on sequence alignments to empty insertion sites. The Region Finder Module enables the comparative analysis and genotyping of elements, fragmented elements, and insertion sites across multiple individuals by grouping sequences into homologous genomic regions.

### Gene Finder Module

The Gene Finder Module, which consists of the commands “starfish annotate”, “starfish consolidate” and “starfish sketch”, facilitates the *de novo* prediction of protein-coding sequences of interest in a given genome assembly. Our rationale for using *de novo* gene prediction is that many *Starship*-associated genes, including *Starship* YRs, are poorly annotated or not annotated at all using the default settings of popular annotation pipelines, thus complicating the use of homology-based sequence searches (45, 46). Targeted gene prediction is first executed using the command “starfish annotate”, which runs the program Metaeuk easy-predict using an input file containing amino acid sequences of interest (38, 45, 47). It then runs the program hmmsearch with a profile HMM of a domain found in the amino acid sequences of interest to filter out sequences that do not contain a high-scoring hit to the domain (47). For example, to annotate *Starship* YRs, users execute “starfish annotate” using as input a file of known *Starship* YR amino acid sequences and a profile HMM built from reference *Starship* YR sequences. If the user additionally provides a GFF file of existing gene features, the *de novo* predicted sequences are named according to any existing gene whose coordinates overlap it to facilitate integration with existing gene annotations. However, all *de novo* predicted genes retain their *de novo* predicted coordinates. “starfish annotate” outputs a GFF file with gene feature coordinates as well as a file containing predicted amino acid sequences. The command “starfish consolidate” may then be used to merge the output with existing GFF and predicted amino acid sequence files, if they exist. Amino acid sequence files and profile HMMs of *Starship*-associated genes are distributed and updated as part of the starfish package, and include *Starship* YRs and the auxiliary genes DUF3723s, ferric reductases (FREs), patatin-like phosphatases (PLPs), and NOD-like receptors (NLRs) (35).

After the gene prediction step, a GFF containing the coordinates of *Starship* YR genes is passed as input to “starfish sketch”, which assigns neighboring genes to gene neighborhoods. Genes within a user-defined base pair distance of each other are placed into the same neighborhood and each neighborhood is assigned a unique identifier that will eventually become the identifier of a mobile element, if detected. The neighborhood organization step is taken to avoid false positive predictions and overlapping element boundaries when *Starship* YR genes are found in close proximity to each other, as the search for element boundaries can be restricted to the upstream- and downstream-most *Starship* YR within a given neighborhood (see next section). “starfish sketch” outputs a BED file with the coordinates of each *Starship* YR for each gene neighborhood.

### Element Finder Module

The Element Finder Module consists of the commands “starfish insert”, “starfish flank” and “starfish summarize”. Together, these commands execute the *de novo* prediction of key mobile element features, including element boundaries, direct repeats (DRs), terminal inverted repeats (TIRs) and empty site coordinates (Figure 2). At each major step, checkpoint files are printed that enable users to restart a particular command if interrupted, or to add new assemblies to an existing analysis.

**Figure 2:**
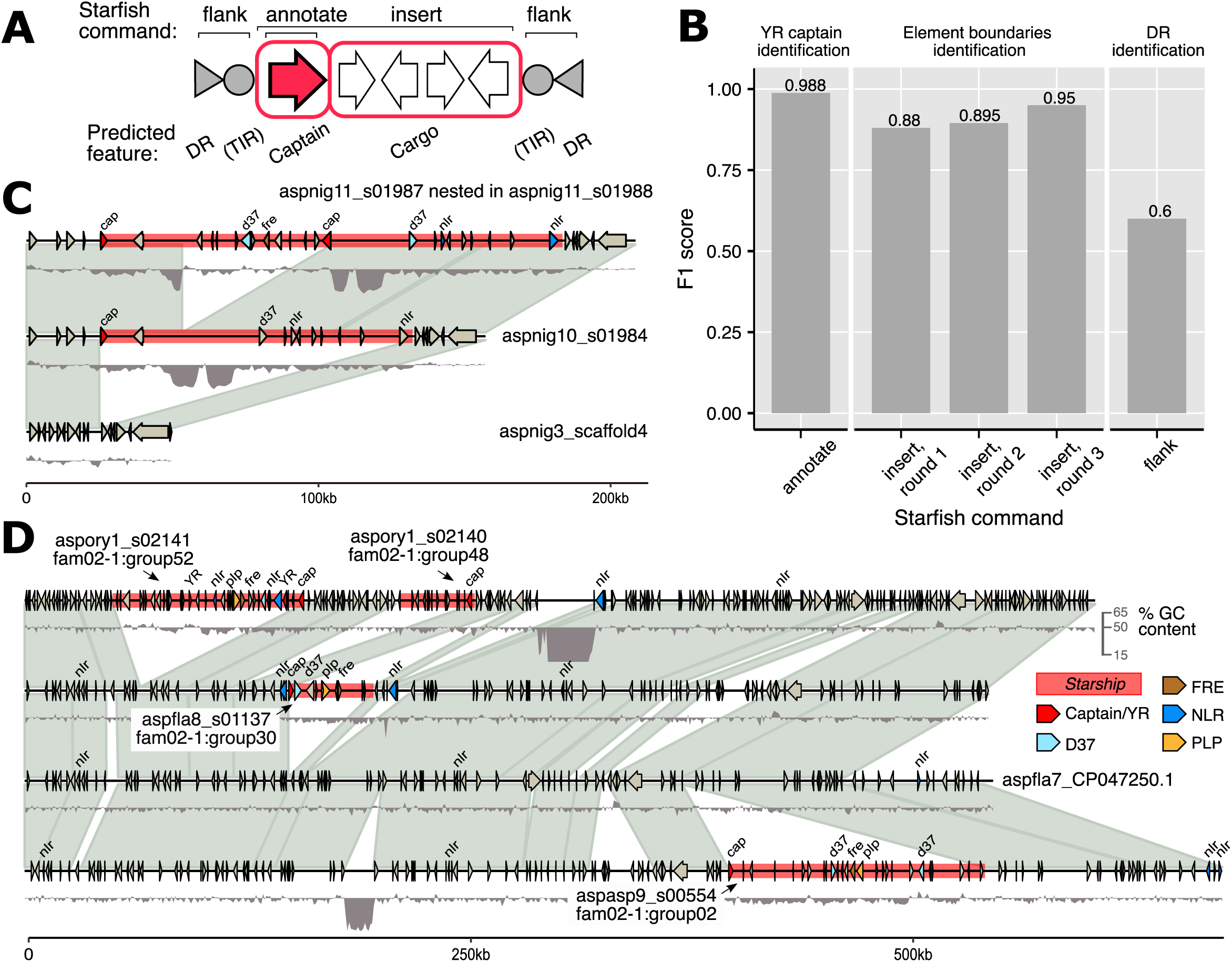
starfish recovers known *Starship* tyrosine recombinases (YRs) and full length *Starship* elements with high recall and precision. A) A simplified schematic of a typical *Starship* element, with features predicted by specific starfish commands indicated. B) A bar graph summarizing the F1 score, the harmonic mean of recall and precision, of three starfish commands for identifying *Starship* YRs, *Starship* element coordinates and direct repeats (DRs), as benchmarked against a database of 42 manually curated elements. The “starfish insert” command was run three iterative times with different parameters (Methods). C) starfish resolves nested element insertions. D) starfish enables comparative analysis and genotyping of complex regions involving multiple independent *Starship* insertions across species. TIR: terminal inverted repeats.

First, “starfish insert” takes as input a BED file of *Starship* YR neighborhoods (output by the Gene Finder module) along with associated genome assemblies. It then conducts sequential BLASTn alignments to find the sequences flanking an element insertion in one individual using the sequences flanking the corresponding empty site in another individual, leveraging the fact that the upstream boundary of all known *Starships* is typically located within 7 kbp (and usually within 2 kbp) of the captain’s transcriptional start site (35, 38).

The upstream sequence “A” of captain “A” from element “A” on contig “A” (controlled with the parameter --upstream) is used as a query to search each assembly for its highest scoring hit “B” (controlled with the --pid and --hsp parameters). The downstream sequence “B” of hit “B”(controlled with the parameter --downstream) is then used as a query to search contig “A” for a high scoring hit “C”. If hit “C” is located downstream of captain “A”, then hit “B” is aligned to its downstream sequence “B” to ensure that they do not match, as a repetitive element could result in a false positive prediction (controlled with the parameter --flankcov). The contiguous sequence of hit “B” plus its downstream sequence “B” is then re-aligned to the region spanning from upstream sequence “A” to hit “C” in order to refine the alignment coordinates, which used to define candidate elements boundaries and boundaries for the associated empty site of the newly defined element “A”. The upstream sequences of the upstream- and downstream-most *Starship* YR within each neighborhood are used to search all provided genome assemblies, resulting in multiple candidate boundaries and candidate insertion sites associated with each *Starship* YR. To avoid false positives that we observed to occur during testing when inversion breakpoints were located within an element, candidate boundaries are then filtered by aligning the sequence of element “A” to contig “B” with nucmer (48) to ensure that its sequence is absent from contig “B” above a given threshold (controlled with the --insertcov parameter). “starfish insert” then outputs a BED file with all candidate element boundaries and *Starship* YRs. Starfish prioritizes user input and flexibility. While all commands have recommended default values, users are encouraged to explore different parameters that may be better suited for their own data. To facilitate this, the commands “annotate” and “insert” may each be run in an iterative fashion enabling users to explore this parameter space through chaining together commands.

Next, the command “starfish flank” is used to search the upstream and downstream sequence around the candidate element boundaries for direct repeats (DRs) and terminal inverted repeats (TIRs) using a version of CNEFinder modified to annotate short repetitive sequences (49). If repeats are found, new boundary coordinates are output reflecting the coordinates of those repetitive sequences. “starfish flank” then outputs a BED file with the coordinates of candidate DRs and TIRs and new boundaries associated with each *Starship* YR.

Finally, the command “starfish summarize” consolidates the results of “starfish insert” and “starfish flank”, primarily by selecting the ‘best’ pair of boundaries that would maximize the length of the candidate element and the length of the predicted DRs, within a given threshold. During testing, longer DRs were associated with higher quality alignments between the element flanking sequence and the empty site flanking sequence, and hence, higher quality predictions of element boundaries. If two candidate *Starship* YRs are located within a neighborhood, one at the 5’ end and one at the 3’ end, at most 2 pairs of candidate boundaries would be kept per neighborhood (one per *Starship* YR) and would require manual inspection by users to select the most appropriate one. “starfish summarize” outputs a BED file with the boundary coordinates and *Starship* YR of each annotated element, and two text files ending in .feat and .stats that respectively contain metadata associated with each element and each empty site. Each element is assigned to either the ‘insert’ or ‘flank’ category, which reflect the measure of confidence in its predicted boundary. ‘Insert’ elements have boundaries informed only by “starfish insert”, while ‘flank’ elements have boundaries informed by both “starfish insert” and “starfish flank”. Existing gene annotations may also be provided to “starfish summarize” to populate the BED file with those annotations.

### Region Finder Module

The Region Finder module consists of the command “starfish dereplicate”. The command genotypes empty and occupied element insertion sites across multiple individuals and groups these loci into homologous genomic regions based on conserved combinations of orthologous genes groups (OGs) flanking the insertion sites. Practically speaking, it is useful for identifying independent segregating insertions of the same element. Key parameters used to control the search include --flanking, the number of OGs up- and downstream of the insertion site that are used to determine region homology; --mismatching, the max number of OG mismatches when grouping into regions; --distance, the max length of up- and downstream sequence to search for flanking OGs; --restrict, which ignores any OG found in at least one predicted element when determining region homology. Users can optionally provide a GFF file containing the coordinates of repetitive sequences (e.g., small TEs) so that OGs that overlap with these sequences can be ignored as well.

”starfish dereplicate” assigns a haplotype label to each sequence within each genomic region: element haplotypes, for sequences containing at least 1 element; empty haplotype, for sequences where the flanking OGs are adjacent to each other; and fragmented haplotype, for sequences where the flanking OGs have other intervening OGs between them. The command outputs several results files: regions.txt which contains coordinates and metadata associated with each sequence in each region; summary.txt, which provides relevant statistics for each region; and dereplicated.txt, which provides a list of representative elements per region.

### Benchmarking data

We assembled a database of 42 manually curated elements, many of which were previously published, from publicly available genomes to evaluate the performance of starfish (Table S1) (35, 38). We calculated recall (True Positives / True Positives + False Negatives), precision (True Positives / True Positives + False Positives) and the F1 score, which is the harmonic mean of recall and precision, for the commands “starfish annotate”, “starfish insert” and “starfish flank”. We used default settings, except for “starfish insert”, where we iteratively chained together successive commands with modified thresholds for --upstream, --downstream and --pid, which respectively modify the length of sequence upstream of the *Starship* YR used in sequence searches; the length of sequence downstream from the flank of the empty site; the minimum percent identity of BLAST alignments.

### Sequence Data

We used the Mycotools software suite to identify, download and format a database of 2899 publicly available fungal genome assemblies (Table S2) (50). We used a tiered sampling approach to sample for both breadth of taxonomic diversity across the fungal tree of life as well as depth within specific fungal genera, including *Alternaria* (Dothideomycetes), *Aspergillus* (Eurotiomycetes), *Fusarium* (Sordariomycetes) and *Verticillium* (Sordariomycetes), that have previously been shown to carry *Starship* elements (35).

### Reference YR sequences and phylogeny

We applied starfish’s Gene Finder module to the genome database to comprehensively annotate *Starship* YRs across the fungal tree of life. We *de novo* predicted a total of 10, 771 YR gene sequences across the genome database (Supporting data; Table S3). To filter out gene fragments and pseudogenes, we removed all sequences <25% of the median predicted amino acid sequence length of 812 residues, obtaining a set of 8369 gene sequences. We then applied best-practice filtering steps based on the recommendations of the UniProt Reference Cluster database to obtain a dataset of representative sequences spanning the full diversity of fungal YRs (Table S4) (51). We used MMseqs easy-clust to cluster sequences by 90% sequence identity to and 80% overlap with the longest seed sequence of the cluster (--alignment-mode 3 --min-seq-id 0.9 --cov-mode 0 -c 0.80 --cluster-reassign) (52). We took the resulting set of representative sequences and again used MMseqs easy-clust, this time clustering sequences by 50% sequence identity and 80% overlap (--alignment-mode 3 --min-seq-id 0.5 --cov-mode 0 -c 0.80 --cluster-reassign). We combined this set of genes with 25 previously annotated captains and 3 fungal DNA CryptonF sequences and aligned them using the E-INS-I method implemented in MAFFT, which is recommended for sequences where several conserved motifs are embedded in long unalignable regions (35, 53). We then trimmed the alignment using clipkit, removing any columns where ≥90% of sequences had a gap (--mode kpic-gappy --gaps 0.90; Supporting data) (54).

We extracted the aligned and trimmed sequence of HhpA, the only functionally validated *Starship* YR, and aligned it to the the YR references cre and XerD (UniProt accessions P06956 and P0A8P8) using the HHpred server for remote protein homology and structure detection (38, 55). The resulting alignments were manually adjusted according to curated alignments of the core-binding (CB) domain and the catalytic (CAT) domain from phage and bacterial YRs (56). The CB and CAT domains are the two main functional domains of YRs, where CB binds the recombination site and CAT catalyzes the cleavage and joining reactions (56). The sequences of these domains, which cover Patches 1-3 and Boxes 1-2 as well as the 6 canonical active sites whose mutagenesis abolishes YR activity, have successfully been used to characterize the phylogenetic history of bacterial YRs (46). We therefore annotated sites in the fungal YR alignment homologous to the CB and CAT domains using the coordinates of the HhpA-cre/XerD alignments as a guide (Table S5). We removed any sequence missing a residue in >1 of the columns corresponding to the 6 known active sites in order to filter out possible pseudogenes, resulting in a final alignment containing 1, 222 representative *Starship* YR sequences and 3 fungal DNA CryptonF sequences.

We extracted the first 512 columns of the reference *Starship* YR alignment, which correspond to the CB and CAT domains plus 50 flanking sites, to build the reference YR phylogeny (Supporting data). We built three maximum likelihood trees using IQTREE v2.0 with automated model selection, 1000 SH-ALRT iterations and 1000 ultrafast rapid bootstraps (-m MFP -alrt 1000 -B 1000), and selected the one with the highest likelihood (Supporting data). We rooted the tree with fungal Crypton DNA transposons, which have previously been identified as relatives to *Starship* YRs (35).

### Superfamily, clade and family classification

We developed a framework that describes *Starship* diversity in accordance with naming conventions proposed by Storer et al. and in an effort to create an operationally useful taxonomy that reflects the multi-partite architecture of CCEs (23, 57). *Starships* belong to the following Dfam lineage: Interspersed repeat; Transposable element; Class II DNA transposition; Tyrosine recombinase; Starship (57). For convenience and compatibility with other classification schema, we consider *Starships* as a “Superfamily” rank equivalent to “Cryptons” within the hierarchy proposed by Wicker et al., and as a “Subclass” within class II YR transposons equivalent to “Cryptons” in the hierarchy proposed by Wells and Feschotte (21, 58).

We then parsed the rooted reference YR tree to identify strongly supported monophyletic clades of *Starship* YRs and develop a more refined taxonomy of this superfamily. Our framework is based on a phylogenetic hierarchy because other mobile element classification schemes based on sequence similarity (e.g., the 80-80-80 rule) do not always result in monophyletic groups, complicating comparative evolutionary analyses (1). After collapsing poorly supported bipartitions with <80% SH-ALRT and <95% ultrafast bootstrap support, we identified three clades present at the polytomy at the root of the tree. We then traversed the nodes that descend from each clade’s ancestor in a stepwise fashion and defined the first encountered clade with ≥20 members and strong support as a *Starship* “family”, until no more families could be defined in this way.

### Navis and haplotype classification

Individual *Starship* elements often have dissimilar features compared with other elements of the same family, indicating that additional ranks below the family level are necessary to fully conceptualize *Starship* diversity. For example, many elements from the same family have different DRs and TIRs, have highly dissimilar captains YRs, and share little to no cargo in common. We therefore developed a classification scheme that incorporates two additional ranks below the family level: one based on the relatedness between *Starship* YR sequences, and one based on sequence similarity across the entire length of the element.

First, within each family, we define “naves” (“navis”, singular; latin for “ship”) based on relationships between *Starship* YRs. We suggest using ortholog relationships between *Starship* YRs as the basis for quantifying these differences, where each ortholog group is a navis assigned a unique name. For example, *Starships Mithridate* and *Aristaeus* both belong to the Hephaestus-family, but their captains differ such that they are typically assigned to different ortholog groups; hence, their separate names are justified and we distinguish them at the navis level (38). We suggest that ortholog relationships be either obtained through ortholog group analysis of *Starship* YR sequences or through the delineation of monophyletic clades on a *Starship* YR phylogeny, two common approaches for annotating gene orthogroups (52, 59).

To then distinguish between *Starships* belonging to the same navis but that carry different cargo, we suggest elements be further defined by “haplotypes” based on comparisons of nucleotide sequence or k-mer similarity scores calculated across the entire length of the element. For example, *Mithridate haplotype1* and *Mithridate haplotype2* are two elements found in *Aspergillus nidulans* and *Penicillium chrysogenum*, respectively, whose captains are highly similar and closely related. However, while they share some cargo in common, including a cluster for formaldehyde detoxification, they differ in total sequence content across much of their length (38). Similar to how gene ortholog groups are frequently determined through a clustering analysis of pairwise E-values using the Markov Cluster (MCL) algorithm (59, 60), pairwise comparisons of nucleotide sequence identity or k-mer similarity scores may also be used as input to a MCL analysis to determine element haplotypes. We provide the commands “starfish sim” and “starfish group” to facilitate the derivation of haplotypes in this way, where the MCL algorithm is applied using the generally accepted default values for a gene ortholog group analysis.

Two elements can thus be said to be copies of each other if they belong to the same family, navis and haplotype. We suggest that elements be named only if insertions of the same copy are found in at least two independent genomic locations within or between species. We refrain from providing exact quantitative thresholds as, similar to species definitions, we believe it is best to be flexible and provide individual *Starships* with unique names and haplotypes when it befits the communication of results. For example, users may instead prefer to define haplotypes based on some other logically consistent criteria, such as the 80-80-80 rule that considers two TEs to be within the same family if they are at least 80 base pairs long and share at least 80% sequence identity over 80% of their length (58). With this in mind, we note that the same copy of an element may undergo changes to its sequence post-insertion and that copies of the same *Starship* may diverge solely due to repeat-induced point mutation (RIP), a fungal genomic defense mechanism, or the nesting of other resident TEs. Depending on the degree of divergence, these copies may be assigned different haplotypes despite being inserted at the same locus, although in the case of RIP where there is evidence that the mutagenized element may no longer be active, we recommend referring to the non-functional copy as a “derelict” or “degraded”.

### *Starship* YR enrichment

We used the size-filtered set of 8369 YR genes and 1422 genomes from 825 species belonging to 10 taxonomic classes that had at least 2 species with an annotated YR sequence to test whether specific YR families are enriched in fungal species from specific taxonomic classes. Using only YRs that were assigned to a specific family, we conducted Fisher’s exact tests for enrichment for each YR family in each fungal taxonomic class using the fisher.test function in base R (alternative = greater), where counts in each contingency table represented the number of species with or without a YR gene from a specific family (Table S6) (61). We applied the Benjamini-Hochberg false discovery rate correction for multiple testing using the function p.adjust in R (61). We calculated the prevalence of each YR family in taxonomic classes of interest by dividing the number of species in the focal taxonomic class with at least 1 member in the focal YR family by the total number of species in the focal taxonomic class.

### *Starship* element and insertion site annotation

We used the Element and Region Finder modules to predict *Starship* elements and their insertion sites from the unfiltered dataset of 10, 771 YRs. We used the unfiltered YR dataset because our primary objective at this step was to recover full length *Starship* elements, which could be associated with YRs of any length. We removed all predicted elements <15 kb in length in order to remove likely false positives. We manually inspected the insertion site alignments of all elements with the command “starfish pair-viz”, and removed any element with a low confidence score that could not be placed into a genomic region by the command “starfish dereplicate”, resulting in a final list of 597 elements in 313 genomes from 144 species (Table S7, Table S8, Supporting data). We removed 38 elements that have >1 candidate captain gene, resulting in a list of 559 high confidence elements used for downstream analyses. We grouped these elements into naves based on the orthologous gene group of their *Starship* YR and into haplotypes based on k-mer similarity over the entire element length calculated with sourmash and MCL through the commands “starfish sim” and “starfish group” (Table S9) (60, 62). We calculated pairwise Jaccard similarity of cargo gene content for each pair of *Starships* using EggNOG ortholog annotation data (Number of orthologs present in both *Starships* / Number of orthologs present uniquely in each *Starship*), and calculated the patristic distance between each captain using the patristic_distances function in phykit on the captain YR phylogeny (Table S10, Supporting data) (63).

We selected one representative copy per family-navis-haplotype combination, each of which we consider to be a distinct *Starship* element. Based on visual inspection of a combined captain phylogenetic tree, 8 captains were manually reassigned to different families, and removed from the reference set, resulting in a final list of 348 representative *Starships* from 126 species (Table S11). We identified 222 unique and non-overlapping insertion sites ≤30bp in length (and therefore, likely to reflect a true insertion site) that are associated with the representative set of elements (Table S12). 143 representative elements that could unambiguously be placed into a genomic region by “starfish dereplicate” were selected and manually annotated to validate starfish predictions (Table S13).

### Functional annotations

We annotated the predicted functions of all *Starship*-associated genes using eggNOG-mapper v2 (--sensmode very-sensitive --tax_scope Fungi) (64). All *Starship-*associated gene sequences were grouped into orthologous gene groups using MMseqs easy-clust (--min-seq-id 0.5 -c 0.25 --alignment-mode 3 --cov-mode 0 --cluster-reassign) (52).

We annotated non-protein coding RNA features overlapping with the *Starship* insertion sites using the Rfam database v14.9 in conjunction with Infernal cmsearch (65, 66); using BLASTn in conjunction with a database of fungal 5S rDNA sequences (http://combio.pl/rrna/taxId/4751/; last accessed 03/01/2023) (67) as well as the software tRNAscan (68). We considered an insertion site to contain a rDNA sequence if at least 75% of the rDNA sequence was present. We calculated the AT content of each insertion site taking into account the 100 bp flanking the site. We searched for overlap between insertion sites and predicted gene annotations using bedtools intersect (69).

### Data visualization

Nucleotide alignments between elements and between elements and insertion sites were visualized with circos and gggenomes (Supporting data) (43, 44). Phylogenetic trees were visualized using ete3 and ggtree (70, 71). Dotplots were generated using numcer from the mummer 4.0 package with options -b 1000 -c 25 and -l 10 (48). All other graphs were generated with ggplot2 (72).

## Results

### Starfish recovers known elements with high precision and accuracy

The basic architecture of a *Starship* element consists of three to four features: the “captain”, the “cargo”, direct repeats (DRs) and occasionally, terminal inverted repeats (TIRs) (Figure 2A). Captains are DUF3435-domain containing tyrosine recombinases (YRs) encoded at the 5’ end of the element that are necessary and sufficient for *Starship* transposition (38), while the cargo is all mobilized sequence outside of the captain’s open reading frame (35–37). DRs and TIRs (if present) are located at the 5’ and 3’ terminal flanks of the elements and presumably play a role in excision and integration, although this has not yet been demonstrated. Many *Starships* carry “auxiliary genes” belonging to four gene families that were previously shown to have strong associations with *Starship* elements: DUF3723s, FREs, PLPs, and NLRs (35)

We assembled a database of 42 elements with these manually curated features to assess the performance of starfish (Methods, Table S1, Supporting data). We evaluated three commands: “starfish annotate”, which *de novo* annotates YRs and auxiliary genes; “starfish insert”, which determines the coordinates of element boundaries; and “starfish flank”, which identifies DR and TIR sequences at the element boundaries. Starfish recovered the key features of these benchmark elements with a >95% F1 score, i.e., the harmonic mean of recall and precision (Figure 2B). “starfish annotate” correctly predicted 41/42 captain sequences from the benchmarking dataset (Recall = 97.6%; Precision = 100%; F1 score = 98.8%). The single captain that was not predicted is from *Starship Wallaby* and is likely a pseudogene, given the large number of premature stop codons within its putative coding region (located from 6, 806, 200 - 6, 808, 937 on contig HG792016.1).

By then iteratively chaining together separate “starfish insert” commands with different filtering thresholds (Methods), starfish correctly predicted the boundaries within 1 kb of 33/42 elements after round 1 (Recall = 78.6%; Precision = 100%; F1 score = 88%), 34/42 elements after round 2 (Recall = 81%; Precision = 100%; F1 score = 89.5%) and 38/42 elements after round 3 (Recall = 92.7%; Precision = 97.4%; F1 score = 95%). The 1 element with a false positive prediction had its upstream boundary incorrectly predicted by starfish to be 7, 691 bp away from the manually annotated boundary, likely due to the presence of a repetitive sequence. The 3 elements that did not have any boundary predictions either did not have a captain predicted as in the case of *Wallaby* (captain identification is a prerequisite to boundary searching) or do not have clean insertions into an empty site (their boundaries were originally obtained through alignment to a reference element) (35). These exceptions lead us to conclude that starfish will recover the vast majority of elements that have identifiable captains and clean insertion sites.

DR prediction proved more difficult, with “starfish flank” correctly predicting DRs for only 18/42 elements (Recall = 60%; Precision = 60%; F1 score = 60%). We suspect this is due to practical challenges associated with automating the detection of short nucleotide repeats of unknown length over relatively large genomic regions. Serendipitously, while annotating the benchmarking dataset, starfish identified 5 previously unknown elements also present on the benchmarking contigs which the authors had missed during the manual curation process, underscoring the benefits of a fully automated algorithm in lieu of manual curation (Table S14). We additionally found that starfish is capable of resolving complicated insertion events, including nested and tandem insertions, after evaluating *de novo* predicted elements from a large genome database (see below; Figure 2C).

### Extant *Starship* diversity is partitioned into 11 major families

We used phylogenetic relationships among *Starship* YRs, the integrative components of *Starship* elements, to define 11 strongly supported and monophyletic “families” that together make up the *Starship* superfamily (Methods; Figure 3). The 11 families group together into 3 monophyletic clades, where clade 1 contains 5 families and clades 2 and 3 contain 3. Our phylogenetic analysis is based on alignments of the conserved core binding (CB) and catalytic (CAT) domains, the two functional domains of YR enzymes necessary for recombinase activity that have previously been used to develop classification frameworks for bacterial YRs, from 1, 222 representative YR sequences sampled across 2, 899 fungal genomes (Methods) (46, 56). Phylogenetic approaches for defining mobile element families are strongly recommended because they result in a hierarchically consistent taxonomic framework, thereby avoiding drawbacks of sequence similarity based approaches that may result in paraphyletic and polyphyletic groups being assigned the same name (21, 23, 57, 58, 73, 74).

**Figure 3:**
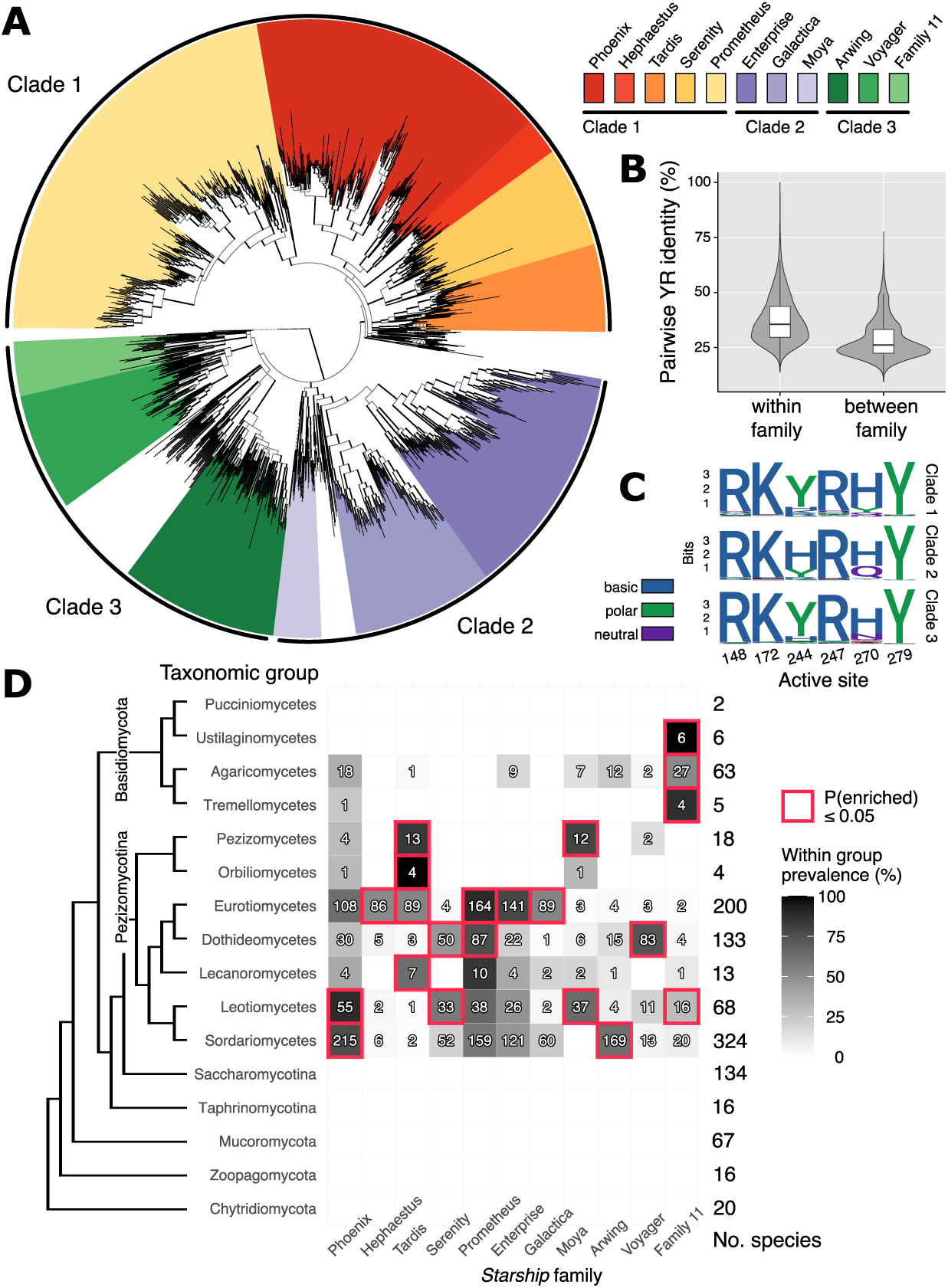
*Starship* tyrosine recombinases (YRs) group into 3 clades composed of 11 families with variable distributions across dikaryotic fungi. A) A maximum-likelihood tree of 1, 222 representative *Starship* YRs built from an alignment of conserved core binding (CB) and catalytic (CAT) domains. The tree is rooted using three fungal CryptonF DNA transposons as the outgroup. All branches with <80% SH-aLRT and <95% UFboot support have been collapsed. Branches are color coded according to *Starship* family. B) Pairwise amino acid sequence identity of all representative YR sequences, calculated from the conserved CB and CAT domain alignment and partitioned according to within and between family comparisons. C) Sequence motifs at each of the 6 active sites present in YR enzymes. Active sites are numbered according to the position of characterized sites in the bacterial YR reference xerD (Uniprot accession P0A8P8). D) To the left is a tree schematic summarizing evolutionary relationships between 11 fungal taxonomic classes in which *Starship* YRs were detected. To the right is a heat map summarizing the prevalence of *Starship* YRs from each *Starship* family in each taxonomic class, where prevalence was calculated as the number of species with at least 1 YR from a given family divided by the total number of examined species in that taxonomic class. Red outlines indicate statistically significant (Fisher’s exact test; P<0.05) enrichment of a particular *Starship* family in a given taxonomic class.

*Starship* YRs are notably diverse at the sequence level, even when only taking the conserved domains into account. The median pairwise amino acid identity within families calculated from the CB and CAT alignment is 35.4% (25th percentile: 29.5%; 75th percentile: 43.6%) while the median amino acid identity between families is 25.8% (25th percentile: 22.3%; 75th percentile: 32.4%) (Figure 3B). However, we identified several conserved residues at known active sites. YRs have a well described catalytic R-K-H-R-Y pentad motif, which is spaced across larger structural regions within the CB and CAT domains (46, 56, 75). *Starship* YR protein sequences are largely conserved at these residues. However, clades 1 and 3 have a YXXR motif in the central residues of the catalytic domain while clade 2 sequences typically maintain the canonical HXXR motif, consistent with previous observations (37). Specific families also differ in the presence and absence of larger indels in conserved regions of the alignment: for example, different families have variably conserved indels across a 19 residue long region in Box 1 and a 2 residue long region in Patch 2 (Figure S1).

Approximately 9% of the representative YR sequences in the phylogeny could not be grouped into a specific family, and remain classified only at the clade level. We suspect this is due in part to the clade-level sequences being shorter than the family-level sequences on average (average length of 464 vs. 588 residues). In particular, clade 3 contains many sequences that were initially grouped into families, but were ultimately determined to be likely pseudogenes given large gaps in the conserved domain alignment indicative of shorter predicted sequences. Following Urquhart et al., we named each family after the first manually curated cargo-carrying element belonging to that family (see below), e.g., the Hephaestus*-*family named after the type element *Starship Hephaestus* (38). All families except Family 11, for which we could not find a cargo carrying *Starship* element associated with a YR sequence in this family, were eventually assigned names in this way (Table 1).

### *Starship* families are variably enriched across taxonomic classes

Each *Starship* family is significantly enriched in at least one fungal taxonomic class (Methods; Figure 3D). Families differ in their enrichment patterns, with some families being enriched in only a single class (e.g., the Hephaestus*-*family) while others are enriched in up to four (e.g., the Tardis-family). *Starships* from multiple families are highly prevalent across the Pezizomycotina. For example, all 11 families are present in at least 1 species from the Eurotiomycetes, Dothideomycetes and Leotiomycetes, and 10 and 8 families are present in at least 1 species in the Sordariomycetes and Lecanoromycetes, respectively. The Pezizomycetes and Orbiliomycetes are exceptions as they generally have few annotated *Starship* YRs. In contrast to the Pezizomycotina, Basidiomycete species generally have few *Starship* YRs and most are missing them all together. No YR genes were found in any of the 11 examined classes outside of the Basidiomycetes and Ascomycetes, and none were found in the Ascomycete sub-phyla Saccharomycotina and Taphrinomycotina (Figure S2, Table S8).

### Starfish predicts new *Starships* carrying diverse cargo

Only a subset of the annotated captain proteins were assigned to cargo-carrying *Starship* elements. While the sequences that could not be assigned to a cargo-carrying element meet all the criteria for being a *Starship* YR from the *Starship* superfamily, it is unclear whether they are “solo” captains that do not transpose additional sequences or whether we simply could not identify the partial or full-length element with which they associate (see Discussion). In total, we identified 348 high-confidence *Starships* segregating at independent genomic regions across 127 species by applying the Element Finder and Region Finder modules to the complete set of *Starship* YR genes (Methods; Table S7, Table S11). The high-confidence elements belong to 9 of the 11 *Starship* families. We searched publicly available genomes, which were not included in our database, in order to validate the 2 remaining families missing a representative *Starship* element. Through manual annotation, we successfully annotated *Starships* from 1 additional family (Methods; Table S13). We could not identify any cargo-carrying *Starship* elements belonging to Family 11; as such, although it is clearly part of the *Starship* superfamily, it remains unnamed and it is not known whether sequences from this family are capable of mobilizing additional cargo.

Median element length ranges from 27, 297-97, 356 bp across the 7 families with ≥10 members (Figure 4A). Across all families, total element length varies 34 fold, with the smallest *Starship* carrying 15, 020 bp and the largest *Starship* carrying 511, 117 bp. We compared the number of shared orthogroups between each pair of high-confidence elements to calculate overall cargo similarity and found that similarity decreases rapidly as *Starship* YRs diverge from each other (Figure 4B). Elements whose captains belong to the same navis (i.e., orthogroup) have an average cargo gene jaccard similarity of 13.4% (Standard deviation = 13.1%) while elements whose captains belong to the same family have an average cargo similarity of 2.5% (Standard deviation = 6.7%). We visualized alignments for three comparisons that illustrate the variety of relationships between captain relatedness and cargo similarity (Figure 4C). These examples demonstrate that captains from the same navis may carry either similar or dissimilar sets of cargo while captains from different naves, although typically carrying dissimilar cargo, have also been found to carry near identical haplotypes.

**Figure 4:**
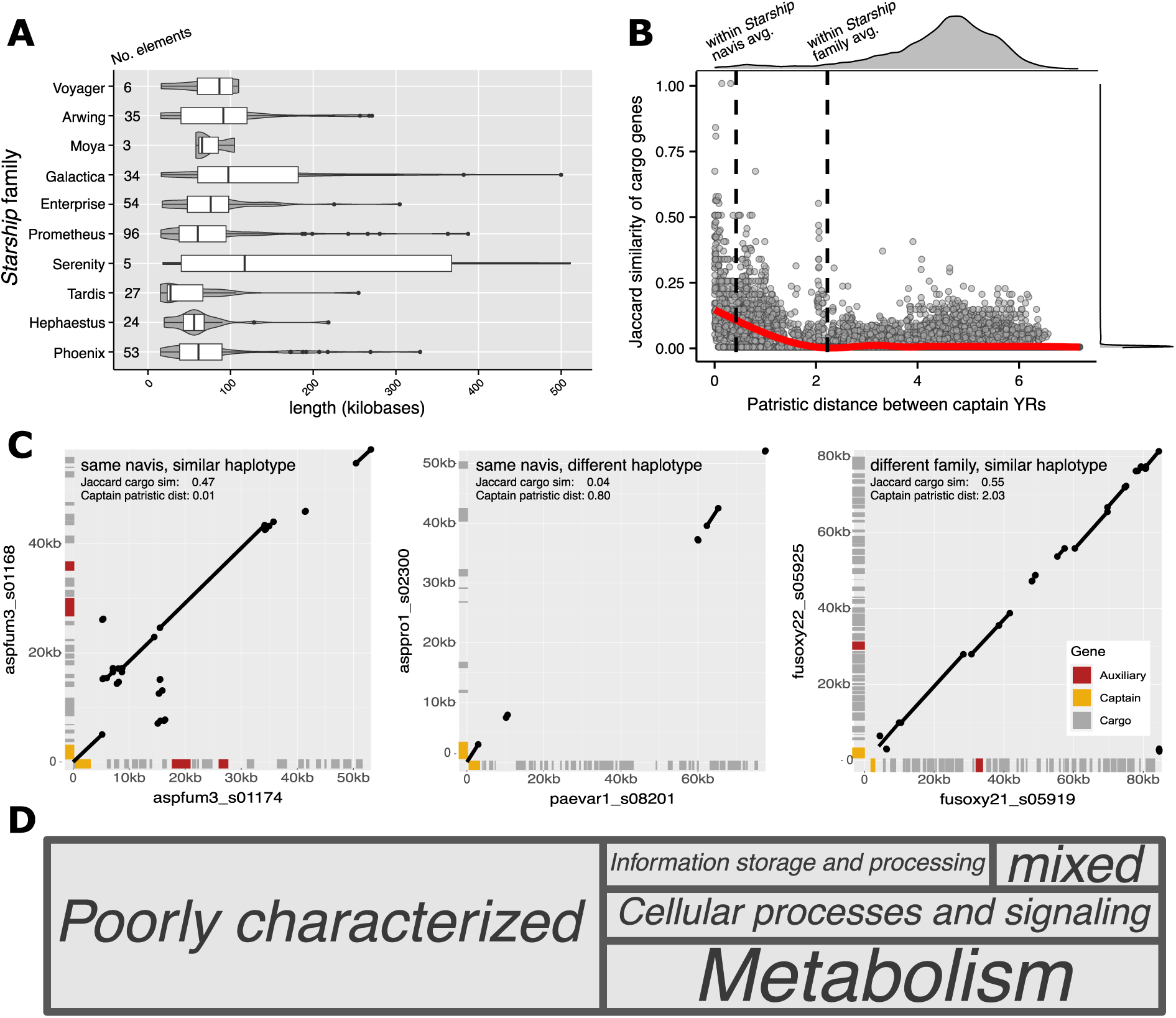
Features of *Starship* cargo. A) Box-and-whisker plots summarizing the length of 348 representative *Starship* elements, organized by *Starship* tyrosine recombinase (YR) family. B) A dot chart summarizing the relationship between the pairwise patristic phylogenetic distance of *Starship* captain YR sequences and the Jaccard similarity of cargo EggNOG orthogroup presence/absence profiles for the 348 representative *Starship* elements. The average within *Starship* navis and average within *Starship* family patristic distances are annotated by vertical dotted lines. A regression line calculated from a general additive model is drawn in red (R^2^ adj. = 0.30). C) Dot-plots visualizing nucmer nucleotide alignments between three pairs of *Starships* of interest that highlight variation in captain YR and cargo sequences. Predicted genes are indicated in the margins of the plots and color coded according to category, where auxiliary genes contain either a DUF3723, PLP, FRE or NLR domain. D) Predicted COG categories assigned to 1909 gene orthogroups carried by 348 representative *Starships*. An additional 2, 541 gene orthogroups (57.1% of total) are not assigned to any COG category and are not shown.

The vast majority of *Starship* cargo has no known predicted function (Figure 4D). Out of the 4450 gene orthogroups carried by the 348 high-confidence elements, 57.1% have no COG annotation, while an additional 21.4% belong to the poorly characterized “General functional prediction only” and “Function unknown” COG categories. The high proportion of unknown cargo is similar to bacterial ICEs, where the majority of cargo genes have no known functions (23). Of the 19.1% remaining orthogroups, 8.9% have a predicted function assigned to the “Metabolism” category, 5.9% are assigned to the “Cellular processes and signaling” category, and 4.3% are assigned to the “Information storage and processing” category. By comparison, the number of predicted genes with annotated functions across different fungal genomes ranges from 33-80%, depending on the method of annotation (76–78).

### *Starships* have repeatedly evolved to target specific genomic niches

We manually annotated 141 *Starship* elements from the reference set in an effort to determine if elements from the same family share representative features (e.g., DRs, TIRs, etc). We first found that elements from the same family appear to share core motifs in their DRs but that these sequences diverge from each other over time (Figure 5A). In some cases, DRs have core motifs that are universally conserved within a family, e.g. the Hephaestus-family with TTAC, while others show no evidence of a conserved pattern or consist of few base pairs, e.g. the Galactica-family (Figure 5A, Table S13). Divergence in DR sequence post-insertion could account for some of the variation we observe, although we note that all elements in this analysis are located at different genomic loci. In nearly all cases, DRs are strand specific, with *Starship* orientation always occuring in the same direction, but intriguingly, the Arwing-family seems to insert in either the forward or reverse direction at a core motif of CCCG/GGGC. Some of the target sites are palindromes, which may help explain this pattern.

**Figure 5:**
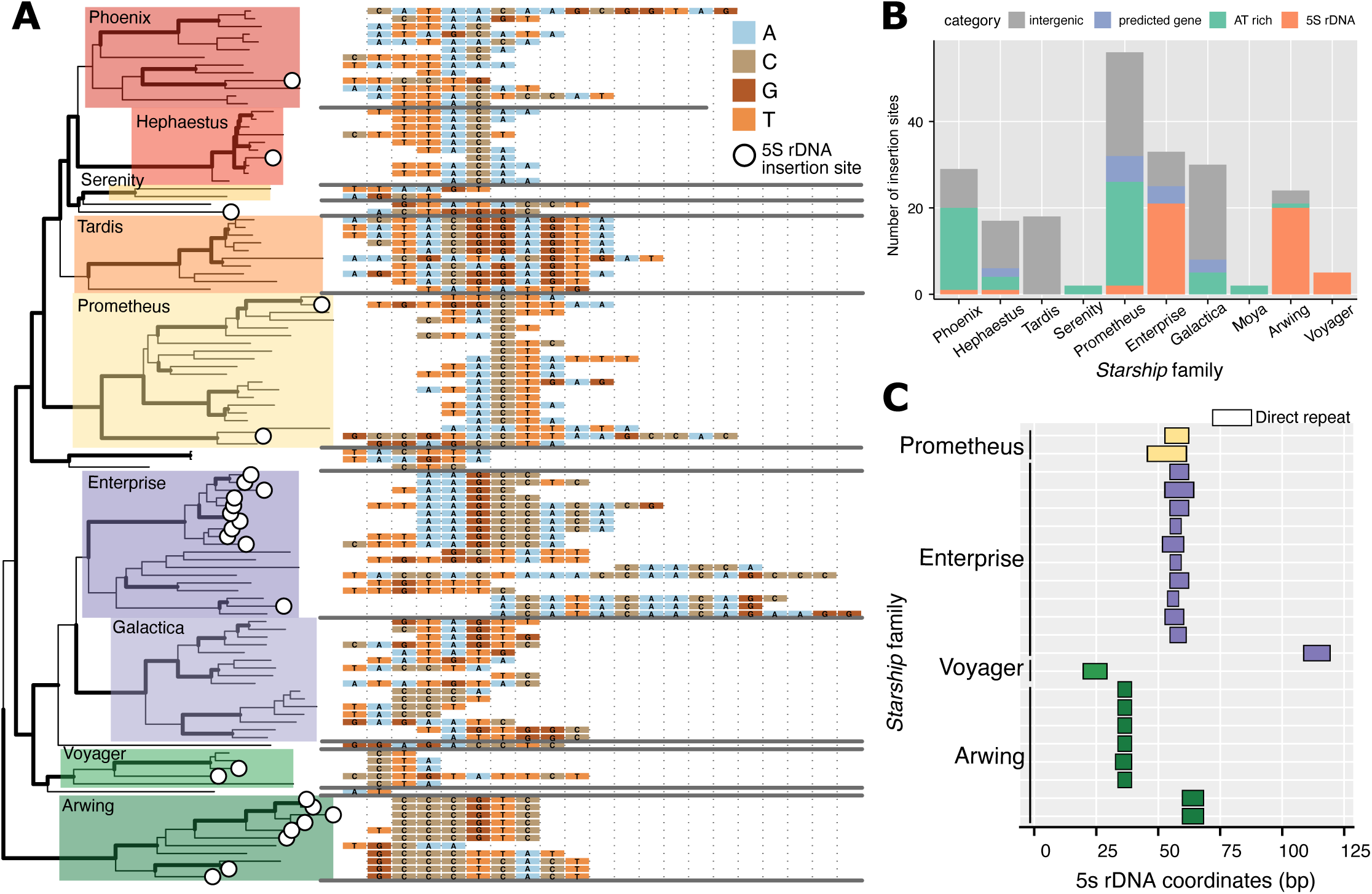
*Starships* target a variety of genomic niches and have repeatedly evolved to insert themselves into 5S rDNA. A) Left: a maximum likelihood phylogeny of 116 *Starship* tyrosine recombinases (YRs) from automated and manually annotated *Starship* elements. Strongly supported branches ≥80 SH-aLRT and ≥95% UFboot support are bolded. Phylogenetic families with ≥2 members have overlaid colored boxes, where coloring is consistent with Figure 1. *Starships* found inserted into 5S rDNA sequence are indicated with a circle. Right: Alignments of manually annotated direct repeats found at the 5’ and 3’ element boundaries. Sequences were manually aligned to each other within each family and not between families. Gray horizontal lines separate sequences from different families. B) A bar chart summarizing the counts of various features of interest found within +/- 100bp of 222 curated insertion sites ≤30bp in length that are associated with representative *Starship* elements. AT rich sequence is defined as ≥60% AT over the +/- 100bp region surrounding the insertion site. C) Coordinates of direct repeats of 23 manual curated *Starships* that are inserted into 5S rDNA, mapped onto the coordinates of matching 5S rDNA sequence and color coded by family. Sequences are vertically arranged in the same order as they appear in the tree in A.

We next examined the nucleotide compositions and genomic context of 222 independent insertion sites associated with the high-confidence elements in an effort to identify trends as to where particular *Starship* families insert themselves in the genome (Methods). We classified 26% of insertion sites as “AT-rich” if the 100 bp region flanking the insertion site was composed of ≥60% A or T nucleotides, which exceeds the genome-wide average of 57.3% AT measured across all genomes included in this analysis (Figure 5B). In many cases we were able to verify that the insertion sites correspond to TEs, suggesting that the *Starships* may either be targeting specific types of sequences or may experience weaker purifying selection at these sites (see Discussion). However, there did not appear to be a correlation with particular TE superfamilies or domains, but this may need to be investigated at finer scales than the family level. As the genome defense mechanism RIP targets duplicated sequences and induces C to T mutations, this observed preference may be related to RIPped sequences specifically (79). Multiple *Starship* families have at least 1 example of an AT-rich insertion site, but some families are more strongly associated than others (e.g., the Phoenix- and Prometheus-families; Figure 5B). Elements from the Serenity-family were particularly difficult to annotate, even manually, as they always appeared in TE-rich regions with poor assembly quality.

We then found that 23% of insertion sites are located within 5S rDNA (Figure 5A, 5B). As with AT-rich sites, 5S rDNA sites are also widely distributed across the *Starship* phylogeny and specific families are more strongly associated with 5S rDNA than others (e.g., the Enterprise-, Arwing- and Voyager-families). Given that direct repeats often reflect the core motif of a transposable element’s true target site (38), we mapped the manually curated direct repeats of 5S rDNA-associated elements onto the coordinates of the 5S rDNA sequence to determine whether different elements may share the same putative target site. In contrast to finding only a single target site, which would be consistent with a single origin of 5S rDNA targeting, we found at least 5 distinct putative target sites associated with different elements, suggesting the ability to target 5S rDNA has independently arisen at least 5 times (Figure 5C). All putative target sites are internal such that an insertion would likely disrupt the function of the transcribed 5S rRNA molecule.

## Discussion

### *Starships* are likely a genetic innovation of the Pezizomycotina

Some of the most striking patterns we observed in *Starship* distributions is the complete absence of *Starship* YRs in lineages outside of the Dikarya, and the general lack of YRs in the Basidiomycota, Saccharomycotina and Taphrinomycotina relative to the Pezizomycotina (Figure 3). Two alternative hypotheses would explain these taxonomic patterns. First, *Starships* may have emerged in the common ancestor of Basidiomycetes and Ascomycetes, followed by subsequent loss in the Saccharomycotina, Taphrinomycotina and multiple lineages of Basidiomycetes. A second hypothesis is that *Starships* arose in the last common ancestor of the Pezizomycotina, and were subsequently horizontally transferred into other lineages outside of this subphylum. Two lines of evidence provide preliminary support for a Pezizomycotina origin, as previously proposed based on the initial description of *Starship* mobile elements (35). First, phylogenetic evidence from numerous systems suggests *Starships* are capable of horizontally transferring between different species (35, 37, 38, 40, 80). Taxonomic constraints on fungal HGT are not known, but while constraints on bacterial HGT are indeed more severe between more distantly related species, they do not preclude horizontal transfer across large phylogenetic distances (81). Indeed, hundreds of genes present in the Basidiomycete genus *Armillaria* are predicted to have been horizontally acquired from Ascomycetes through a yet unidentified mechanism (82). Second, nearly every taxonomic class in the Pezizomycotina is enriched for multiple *Starship* families and has multiple annotated elements. In contrast, few Basidiomycete lineages have *Starship* YRs, and none have *Starship* elements with identifiable boundaries. Additional work is needed to investigate the phylogenetic history of *Starships* in Basidiomycetes to test whether they form a monophyletic group, or whether *Starship* YRs have different common ancestors more closely related to Pezizomycotina sequences, indicative of horizontal transfer.

### Patterns of *Starship* diversity vary across fungal taxa

Differences in *Starship* distributions also manifest over shorter evolutionary distances, suggesting specific demographic forces shape the process of *Starship* diversification. For example, we found that *Starship* families differ in their prevalence and enrichment across fungal taxonomic classes. Consequently, classes often differ in the combinations of *Starship* families found in their genomes (Figure 3). These differences may be due in part to the underrepresented samping of particular lineages, such as the Lecanoromycetes or Orbiliomycetes, which have few publicly available genomes. However, amongst well sampled clades, differences in host range are more likely explained by three non-mutually exclusive hypotheses of lineage expansion and contraction related to niche and neutral theories of biodiversity (7).

First, *Starships* may have adaptations that facilitate their expansion in specific taxa. It remains unclear what these adaptations would consist of, but they likely relate to the evolution of integration site selection through the targeting of particular genetic and epigenetic motifs (see below). For example, MAGGY fungal transposons specifically target genomic sites with H3K9 methylation and several captains from the Enterprise-family have CHROMO domains implicated in heterochromatin binding, suggesting differences in chromatin profiles among species may circumscribe variation in mobile element repertoires (36, 83). Enrichment patterns may alternatively arise through processes acting at the level of the host genome that decrease element frequencies, resulting in family contraction. For example, Urquhart et al. demonstrated that despite their low copy numbers, *Starships* are susceptible to mutagenization and inactivation by RIP (38). Mobile element density is also typically negatively correlated with meiotic recombination rates (84). Fungal lineages with highly active RIP machinery and high recombination rates may therefore be less suitable hosts for larger CCEs (85, 86). Finally, differences in *Starship* distributions may be explained through stochastic processes that underlie element emergence, expansion and contraction. For example, the Hephaestus-family is found primarily in the Eurotiomycetes and its members have among the shortest branch length distance to their most recent common ancestor, suggesting that this family may have a restricted host range simply because it emerged more recently during the course of fungal evolution (Figure 3). Evaluating support for these competing hypotheses and increasing the sampling of underrepresented lineages will help reveal the genetic determinants of *Starship* host range and the mechanisms maintaining and generating CCE diversity more broadly.

### *Starship* niches reflect both specialist and generalist strategies for persistence

*Starship* distributions within fungal genomes suggest they deploy two common strategies for mobile element persistence (87). First, elements from multiple families, including the Hephaestus- and Galactica-families, appear to integrate into randomly dispersed genomic locations, as they are not associated with sites that share any discernible features (Figure 5B). A “generalist” strategy of randomized integration would effectively enable a population of elements to sample a large range of genomic niches to find, by chance, particular sites that maximize their fitness, for example, sites with minimal deleterious costs on their host (9, 36). Although we cannot rule out the possibility that these sites share some as yet unmeasured feature in common, support for the generalist strategy comes from Urquhart et al., who found that *Starship Hephaestus* (the type element of the Hephaestus-family) inserts itself into low specificity TTAC(N_7_)A motifs that are randomly distributed across thousands of possible locations in the genome (38). Indeed, many *Starships* have short DRs (2-6 bp) indicating they may be capable of integrating into thousands of locations, since DRs of *Starships* typically match the core motif of an element’s true target site (38). It is worth noting that the relationship between DRs and target sites is not always straight-forward, and that additional site specific requirements likely exist. For example, the “A” located 7 bp downstream of the core target site motif appears absolutely required for insertion of the *Hephaestus* type element and is not evident from examining the DR, and the true target site of the *Enterprise* type element appears to be a full 10-12 bp palindrome even though the DR is only 5-6 bp (36, 38).

Yet not all insertions appear random, as many elements are inserted into sites that share specific features, notably AT-rich sequence and the 5S rDNA coding region (Figure 5B). On the one hand, the prevalence of these two types of sites may simply reflect survivorship bias, where elements inserted here have higher chances of survival compared with elements inserted elsewhere (9, 88). For example, many fungal genomes have large regions of gene-poor, AT-rich sequence that are typically under relaxed selection and therefore prone to the accumulation of mobile elements (89). Yet if all *Starships* had higher chances of survival in AT-rich or 5S rDNA sites, then we would expect these genomic “safe havens” to be randomly distributed across the *Starship* YR phylogeny. Instead, there are five families biased towards inserting into these particular sites, suggesting that these elements have evolved mechanisms of integration site selection (88).

In the first example of putative site selection, the majority of insertion sites associated with the Phoenix- and Prometheus-families are found in AT rich sequences. Although we could not comprehensively annotate repetitive elements for all genomes, we found through manual curation of target sites from these families that many, though not all, insertions into AT-rich regions are insertions into bonafide TEs. Neither *Starship* family appears to prefer specific classes of TEs, as *Starships* could be found within both Class I and Class II TEs. As individual TEs have their own GC profiles that may differ from the genomic average and the timing of RIP mutagenesis vs. *Starship* insertion is not known, it is difficult to determine if this insertion preference is for nucleotide context, TEs deactivated by genome defense such as RIP and/or methylation, chromatin context, or a combination of these (88, 90). Regardless, the preference does not appear to be tied to specific DRs, as the Hephaestus-family has an AT-rich target site of TTAC(N_7_)A but does not show an enrichment in AT-rich insertions. It is also unlikely to be a result of specific selection regimes in a given organism as many Prometheus- (AT-rich associated) and Hephaestus-family (not AT-rich associated) elements are found within the same genomes, primarily in *Aspergillus*.

Investigating other protein domains beyond the DUF3435 of the Captain proteins may help understand these insertion site preferences. For instance, the *Enterprise* captain contains a CHROMO domain and a Zinc Finger domain implicated in heterochromatin and DNA binding, respectively (36). The Serenity-family is of particular interest for future investigations into insertion site bias. While starfish struggled to annotate full elements in this family, manual efforts revealed that this is likely due to the fact that most of these *Starships* appear to reside in large TE islands, hampering the ability to find clean insertion sites. However, many more high-confidence insertions will need to be found before this trend can be confirmed.

In another example of putative site selection, the majority of insertion sites associated with the Enterprise-, Arwing- and Voyager-families are located within 5S rDNA. Elements from these families have DRs that match specific sequence motifs in the 5S rDNA and are among the longest *Starship* DRs identified to date, ranging from 6-16 bp, indicating these elements likely target 5S rDNA with high specificity. Furthermore, the ability to target 5S rDNA sequence has likely convergently evolved multiple times across the *Starships’* evolutionary history, suggesting the ability to target this site provides a selective advantage (Figure 5C). We therefore consider at a minimum the prevalence of 5S rDNA and perhaps AT- or TE-rich insertion sites indicative of a specialized strategy for maximizing element fitness through the evolution of integration site selection (88).

### Considerations for *Starship* classification

The development of starfish has presented us with a vast diversity of genetic entities to investigate and categorize. Having a classification system is of central importance to the communication of scientific findings, but delineating one is not straightforward, especially for mobile elements (73). With many biological systems, it is difficult to define exact boundaries where divisions should be made, and the *Starships* are particularly recalcitrant to standard methods applied in the eukaryotic TE community due to their size and cargo turnover (57, 58, 74). Our proposed division of 11 families is phylogenetically supported and may require further splitting in the future. Under the current schema, it is possible that members of the same family have low nucleotide identity to one another and may also differ in their target site and/or DR, which is a major challenge in general for coherent TE classification. For example, most *Starships* in the Enterprise*-*family from *Aspergillus* target the 5S rDNA, whereas *Enterprise* itself targets a palindromic sequence. While it may be useful to divide these groups into separate families based on this observation, we refrain from doing so until we have a better understanding of how the important features, like target sites, change over time. If such features diversify too rapidly, then these may be poor characters upon which to define families. If new families are defined in the future, reference elements should be selected along with a monophyletic clade in the *Starship* YR phylogeny. The current framework thus presents a stepping stone towards providing a meaningful basis for describing this new dimension of fungal genomes, leading to a more complete understanding of the processes shaping eukaryotic diversity.

### Usage recommendations and future directions

Starfish establishes a baseline for CCE annotation in eukaryotic genomes by faithfully recovering known *Starships* while greatly expanding our capacity for *de novo* prediction. It thus represents a much needed addition to our collective comparative genomics toolbox and a first step towards making giant CCE prediction routine. Below, we outline several avenues for continuing to improve the workflow in subsequent releases.

The low ratio of *Starship* elements with predicted boundaries to annotated *Starship* YRs (559:10, 771) suggests either that: 1) few *Starship* YRs are associated with cargo sequences; 2) we underestimated starfish’s false negative rate; or 3) our genome database is not well suited for *Starship* identification. Although we suspect that much of this discrepancy is due to the third scenario, some combination of all three are likely responsible. First, not all *Starship* YRs that we identified are expected to be active, and many may be pseudogenes. We plan to implement a filter in future release to screen out pseudogenized *Starship* YRs or those that are part of inactive elements. Second, our benchmarking dataset of 42 elements represents a small fraction of *Starship* and insertion site diversity. As more elements continue to be annotated, we will continue evaluating why starfish fails to recover specific types of elements but not others. For example, we have observed that starfish performs poorly when TEs or low complexity sequences flank an insertion site. Additionally, if a *Starship* is present within a novel TE insertion, starfish may incorporate said TE within the element boundaries. We will evaluate how repeat masking prior to running starfish may help or improve performance in these scenarios. Finally, it is challenging to recover full-length *Starships* from short-read genomes, such as those that make up the database we used here, as many elements frequently exceed the assembly N50 or are located in difficult to assemble regions. More suitable databases will become available in the coming years as population-level long-read sequencing becomes commonplace. These recommendations represent exciting avenues for future development and addressing them will enable us to continue refining our understanding of how mobile elements shape the ecology and evolution of their hosts.

## Supporting information

Supplementary data titles and legends

## Data Availability

All scripts used in this study to generate figures and conduct comparative and statistical analyses, in addition to supporting data such as sequences, alignments and Newick tree files are publicly available from the following Figshare repository (DOI: 10.6084/m9.figshare.24430447). Starfish is freely available as open source code under a GNU Affero General Public License version 3.0 from https://github.com/egluckthaler/starfish.

## Acknowledgments

The authors thank Aileen Berasategui, Johannes Debler and Adrian Forsythe for providing feedback on starfish during beta testing, as well as Tobias Baril, Adrian Forsythe and Andrew Urquhart for providing comments on an early draft of this manuscript. Special thanks to Daniel Croll for providing computational support necessary to conduct these analyses.

