## Supplementary data titles and legends for "Systematic identification of cargo-carrying genetic elements reveals new dimensions of eukaryotic diversity"

Table S1: Benchmarking database metadata and results

Table S2: Genome database metadata

Table S3: Coordinates of 10,771 de novo predicted tyrosine recombinases

Table S4: mmseqs cluster representatives and family metadata for 8369 size-filtered tyrosine recombinases and 25 reference YR sequences

Table S5: Structural alignment coordinates of the representative tyrosine recombinase alignment to tyrosine recombinase reference sequences

Table S6: Contingency tables of tyrosine recombinase families per fungal taxonomic class

Table S7: Coordinates of  Starship elements and their associated features and cargo

Table S8: Number of tyrosine recombinases and Starship elements per genome

Table S9: Starship classification metadata

Table S10: Jaccard similarity in cargo and patristic distance for each pair of Starship captains

Table S11: Metadata for 348 representative Starship elements

Table S12: Metadata for 222 insertion sites <=30bp associated with representative Starship elements

Table S13: Manual curation validation of 143 Starship elements predicted by starfish and 1 additional manually curated element

Figure S1: Visualization of the trimmed *Starship* tyrosine recombinase alignment. To the left is the maximum likelihood tree of 1,222 representative *Starship* YRs built from an alignment of conserved core binding (CB) and catalytic (CAT) domains (as in Figure 1). The tree is rooted using three fungal CryptonF DNA transposons as the outgroup. All branches with <80% SH-aLRT and <95% UFboot support have been collapsed. To the right is the multiple sequence alignment used to produce the tree.

Figure S2: Number of predicted *Starship* tyrosine recombinases (YRs) per fungal taxonomic class. Box-and-whisker plots overlay individual data points with jittered positions. Each data point represents an individual genome.
