## Supplementary figures and images for "Systematic identification of cargo-carrying genetic elements reveals new dimensions of eukaryotic diversity"

### Figure S1

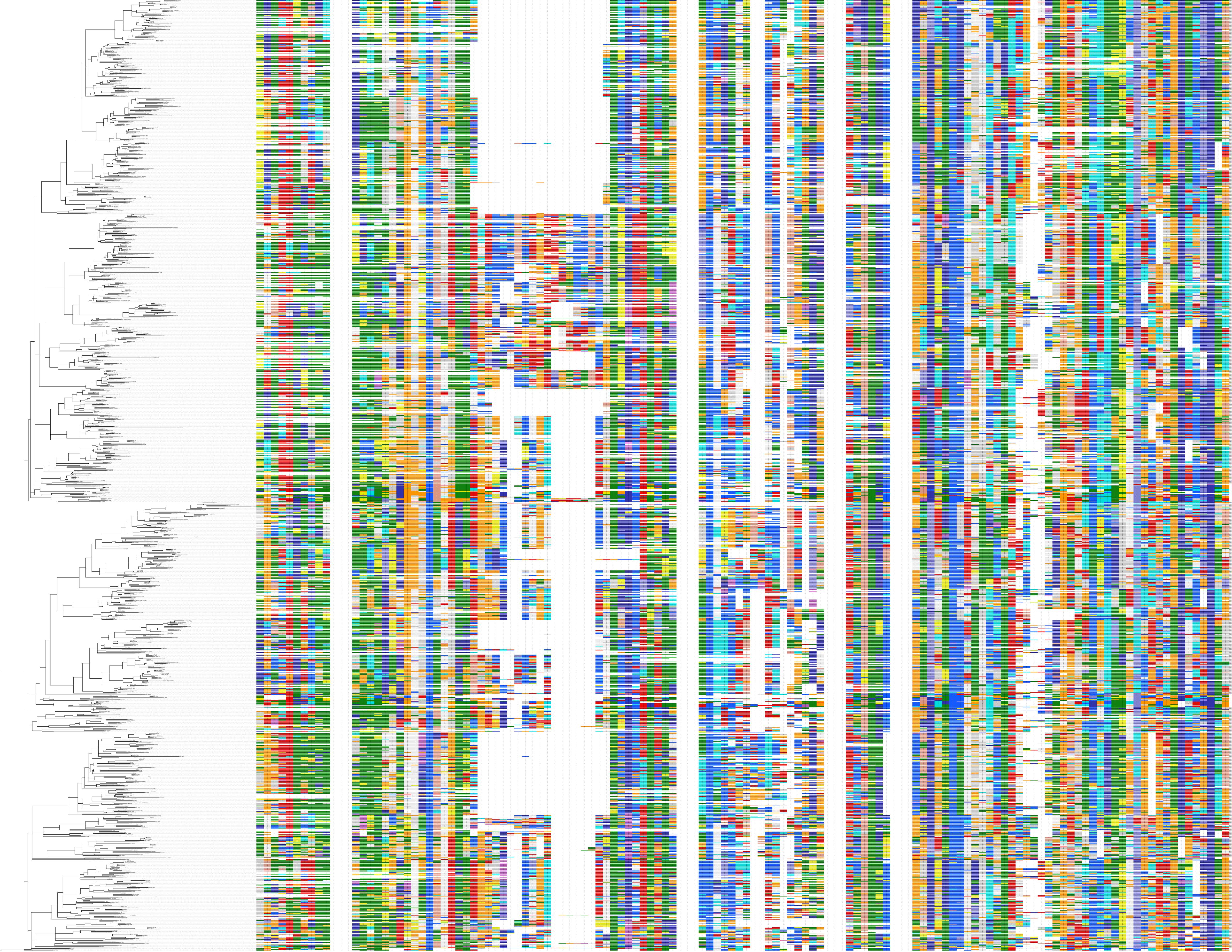

### Figure S2

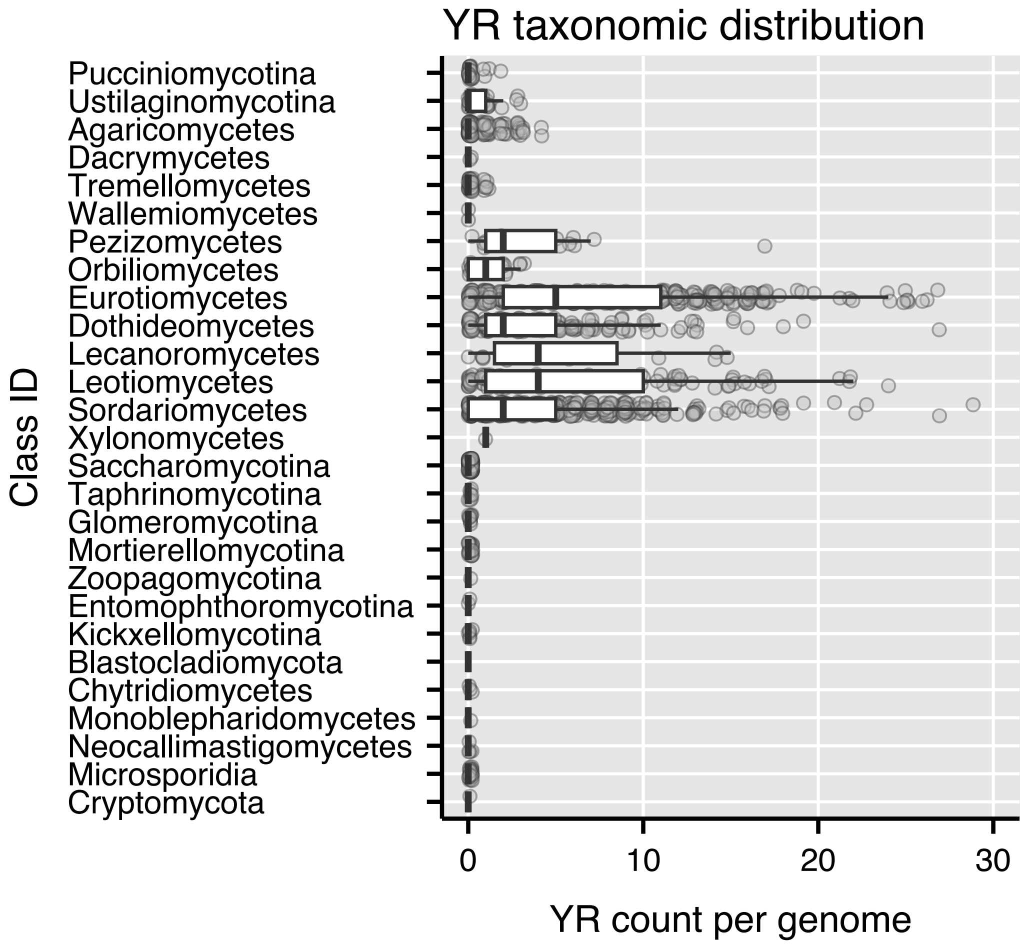
