## Supplementary material for "Systematic identification of cargo-carrying genetic elements reveals new dimensions of eukaryotic diversity": Table 1

| Family Name | Clade | Family Number | Type element described |
| --- | --- | --- | --- |
| Phoenix | 1 | 1 | Gluck-Thaler et al., 2022 |
| Hephaestus | 1 | 2 | Urquhart et al., 2022 |
| Tardis | 1 | 3 | Gluck-Thaler et al., unpublished |
| Serenity | 1 | 4 | This study |
| Prometheus | 1 | 5 | Urquhart et al., unpublished |
| Enterprise | 2 | 6 | Vogan et al., 2021 |
| Galactica | 2 | 7 | Gluck-Thaler et al., 2022 |
| Moya | 2 | 8 | This study |
| Arwing | 3 | 9 | Gluck-Thaler et al., 2022 |
| Voyager | 3 | 10 | Gluck-Thaler et al., 2022 |
| Family 11 | 3 | 11 | n.d. |

Table 1: *Starship* element family metadata
